# Spatial and temporal variation in biting midge (*Culicoides*) abundance in active nest boxes

**DOI:** 10.1101/2025.07.21.665931

**Authors:** Jan-Åke Nilsson, Martin Stjernman

## Abstract

Due to recent and projected global warming, the fitness landscape of organisms can be predicted to change. One of the consequences of climate change is an increased risk of being infected by pathogens. Many of these pathogens are relying on vectors for transmission. As most vectors are ectotherms, with a tight relationship to ambient temperature for e.g. activity, temperature-dependent variation in vector density may be the mechanism explaining increasing parasitemia. In this study, we estimate the abundance of biting midges (*Culicoides spp*), a vector of *Haemoproteus* parasites, by placing sticky traps within nest boxes of breeding blue tits (*Cyanistes caeruleus*). We found that average minimum temperatures positively affect the abundance of midges. Thus, with recent and predicted higher temperatures, the encounter rate between hosts and vectors will increase with a resulting higher parasitemia. Marked differences in spring temperatures will also result in cohort effects, with whole generations of first-year birds being more or less infected by the parasites with yearly effects on recruitment rate and total host densities. Furthermore, we found pronounced temporal and spatial variation within years. This will put increasing emphasis on breeding parents to use cues indicating high vector abundances when determining breeding timing and choice of breeding habitat.

## INTRODUCTION

Recent and future climate change have been predicted to lead to an increase in prevalence and range of vector-borne parasites (Garamszegi 2011; Zamora-Vilchis et al. 2012; Cazelles et al. 2023). Part of this increase could potentially be due to variation in vector abundance, however, to what extent a change in climate affects the activity and life cycle of the vector, resulting in variation in abundance, is poorly known. Most vectors are ectotherms and temperature may hence affect their daily or seasonal activity, reproductive rate as well as life span (Mellor et al. 2000).

The Haemosporidian parasites causing avian malaria, a common and widespread disease among avian populations, are spread by Dipteran vectors. The three most common parasite genera, *Plasmodium*, *Leucocytozoon* and *Haemoproteus*, have each their own Dipteran vector, mainly mosquitoes (*Culicidae*), black flies (*Simuliidae*) and biting midges (*Ceratopogonidae*), respectively (Atkinson & van Riper 1991; Valkiūnas 2005). As Haemosporidian parasites have been shown to induce negative fitness effects on their hosts (e.g. Martínez-de la Puente et al. 2010a; Asghar et al. 2015; Grabow et al. 2025), we would predict that such hosts should try to avoid contact with the vectors as much as possible.

The effect of average ambient temperature on the abundance of black flies and biting midges has been tested but the results are mixed. The abundance of black flies has been found to be negatively associated to ambient temperatures (Martínez-de la Puente et al. 2009a) or not at all (Castaño-Vázquez & Merino 2022) whereas biting midges are sometimes positively affected (Castaño-Vázquez & Merino 2022) or indifferent (Martínez-de la Puente et al. 2009a) to ambient temperature. From a mechanistic point of view, one could predict a positive relation as increasing ambient temperatures increases the level of activity at least among biting midges (Mellor et al. 2000; Tugwell et al. 2021). The effect of wind speed has also produced mixed results with both positive and negative associations with vector abundance (Martínez-de la Puente et al. 2009a; Castaño-Vázquez & Merino2022). These climatic factors have the potential to explain some of the variation between years in vector abundance (Castaño-Vázquez & Merino 2022). Thus, to the extent that these vectors are indeed positively affected by climate change, increased parasitemia in host species may exert a new selective force already now and even more so in the future.

Besides climate driven effects on the abundance of flying vectors, variation in ambient temperatures may change the seasonal activity pattern of the vectors. Such temporal variation within a year could affect the timing of life history stages of the host, for example timing of breeding, to avoid periods with high vector abundance. Biting midges seem to increase in number as the breeding season proceeds and later breeding species encounter a higher number of vectors (Martínez-de la Puente et al. 2009a; Castaño-Vázquez & Merino 2022). Thus, there seems to be a premium in early breeding to avoid the peak of vector abundance.

Spatial variation in vector abundance could also potentially influence breeding site choice in birds. At a landscape level, increased distance from water bodies have been found to reduce the abundance of vectors (Krams et al. 2022). This is parallelled by an increased risk of being infected with *Haemosporidian* parasites with proximity to water (Lachish et al. 2011). However, small-scale variation in vector abundance in seemingly homogenous habitats have received little attention.

At least black flies and biting midges are known to enter cavities and may thereby feed on and potentially transmit *Haemosporidian* parasites to nestlings (Tomás et al. 2008a; Votýpka et al. 2009; Castaño-Vázquez et al. 2021). From a vector point of view, nestlings with undeveloped plumage will facilitate blood sucking and from the parasite point of view, nestlings with a undeveloped immune system will facilitate their establishment within the host.

Here we aim at estimating the degree to which the abundance of *Haemosporidian* vectors will be dependent on ambient temperature by employing traps within nest boxes of breeding blue tits (*Cyanistes caeruleus*). Such a temperature-dependence, resulting in between-year variation of vector abundances, could explain cohort effects on parasite prevalence explaining between-year variation in the density of host populations. Furthermore, we will evaluate the potential effect of temporal (within-year) variation in vector abundance with selection on the hosts to time their breeding to avoid peak vector abundances. If spatial (local) variation in vector abundance is extensive, another behavioural trait that can be predicted to increase in importance is host breeding site selection.

## METHODS

The study, conducted during the spring/summer seasons of 2008 and 2009, is part of a long-term project studying life-history and host-parasite interactions in nest box breeding blue tits (*Cyanistes caeruleus*) in three distinct areas located approximately 20-30 km east of Lund, Sweden. The Revinge area (55.70°N, 13.46°E; elevation: 18-37 m above sea level) is situated in a patchy landscape, consisting of woods and groves of primarily deciduous tree species interspersed with meadows and agricultural land. The areas Öved (55.71°N, 13.61°E; elevation: 43-113 m asl) and Romele (55.62°N, 13.43°E; elevation: 100-129 m asl) are situated in more continuous stretches of deciduous and mixed forests, respectively. In these areas, blue tits and marsh tits breed in nest boxes (inner dimensions: height x width x depth 20 x 8.1 x 9.5 cm; entrance hole diameter: 26 mm, excluding dominant great tits), attached to tree trunks at approximately 1.5 m above ground.

### Field methods

Flying insects were caught within a nest box according to the second method in Tomás et al. (2008a). Specifically, we painted the bottom of plastic petri dishes (diameter: 53 mm) with a thin layer of commercially available baby oil gel (Johnson’s baby oil gel with camomile, Johnson & Johnson, Neuss, Germany). With the help of wire netting (mesh size 1 cm), we placed the petri dishes up-side down in the ceiling inside nest boxes that was either occupied by breeding blue tits (*Cyanistes caeruleus*) or empty (only in 2008). The petri dishes remained in position for two (nestling stage) or four (incubation) days before being removed. During the nestling stages, the old traps were exchanged for new ones after two days, resulting in a total sampling period of approximately 4 days at all stages. All trapped insects were carefully removed, and biting midges (by visual determination) counted and stored in 70 % ethanol for later species identification.

In 2008, we placed traps in 51 nest boxes with incubating blue tit females. The petri dishes were attached to the ceiling of each nest box 4 days before anticipated hatching date (blue tits incubate for a minimum of 12 days in the study area; Nilsson et al. 2008) and removed approximately 4 days later (mean: 3.8 days; range: 2.8 – 4.2 days). This resulted in insect trapping being performed between 15 May and 23 May during the incubation stage. In the same nest boxes, if still active, we placed new petri dishes also during the early and late nestling stage (nestling day 2-6 and 12-16, respectively). On average, we sampled for 4.0 days (N = 42; range: 3.6 – 5.6 days) during the period 22 May to 30 May in the early nestling stage and for 3.9 days (N = 38; range: 3.6 – 4.5 days) during the period 1 June to 9 June for the late nestling stage. During 2008, we placed petri dishes inside 19 unoccupied nest boxes following the same protocol as in occupied boxes and covering the total period 20 May to 10 June.

During the breeding season of 2009, traps were only placed in occupied blue tit nest boxes and only during the late nestling stage, covering the nestling days 12 to 16 during the period 17 May to 9 June. On average, the traps were active for 4.1 days (N = 53; range: 3.8 – 4.9 days).

During the late breeding stage in 2008 and 2009, we calculated mean average, mean maximum and mean minimum ambient temperature from a nearby meteorological station (Swedish Meteorological and Hydrological Institute, station Lund 53430) corresponding to the specific 4-day period when the trap was baited in each nest box.

### Extraction of DNA and sequencing of biting midge samples

We selected 30 biting midges, 15 per sample year, spread throughout the sampling period for molecular species identification. Biting midges were incubated at 56 °C in 100 μl Lysis buffer and 1.5 μl proteinase K for 3 hours. Samples were centrifuged for 10 min at 10000 rpm before the supernatant was removed and put into new tubes of 10 μl NaAc (3M) upon which 220 μl ice cold 95 % ethanol were added. Samples were spun at 10 min at 10000 rpm and the supernatant was removed. An additional washing step was done by adding ice cold 70 % ethanol followed by 5 min centrifugation at 10000 rpm before removing the supernatant. The DNA pellet was dissolved in 25 μl ddH_2_O. The DNA was quantified using a Nanodrop. The amount of DNA extracted ranged from 2.7 to 51.5 ng/μl.

The molecular identification of the biting midges was done by amplifying the COI gene using two different combinations of primers. The primers C1-J-1718 (F) together with C1-N-2191 (R) (Videvall 2013) or C1-J-1718b (F) together with C1-J-1718b (R) (Dallas et al 2003) following the annealing temperatures, precipitation and sequencing protocol described in Videvall et al (2013).

Obtained sequences were aligned and trimmed at the 3′end using Geneious Prime 2019.2.3. Trimmed sequences ended up having 441 bp and were submitted for a megablast, for highly similar sequences, search at GeneBank (Dennis et al 2013) using the standard nucleotide collection. Blast hits with an E-value of 0 and > 99% sequence similarity were assigned to the retrieved species.

### Analyses

All statistical tests were performed with R v.4.5.0 (http://www.R-project.org/). Our count data represents the number of midges caught during an approximate 4-day period. To account for the small variation in exposure time, we always included the specific number of days for each box as a covariate in all models. This specific number of days was never significantly affecting the response variable (all P > 0.24) probably reflecting the general crepuscular and nocturnal lifestyle of biting midges (Mellor et al. 2000). Since all traps were mounted, exchanged and dismounted during daytime, variation in the length of deployment of the traps was due to small variations during the day. To test for variation in no. of caught midges during the 4-day sampling session in relation to the stage of breeding, we included stage (incubation, early and late nestling stage) and site (Revinge, Öved and Romele) as fixed factors as well as the interaction between stage and site using the dataset from 2008. As the same boxes were used during the three stages, box number were included as a random factor. Using data from both 2008 and 2009 but restricted to the late nestling stage, we tested how much of the variation in no. of caught midges during a 4-day sampling session could be explained by year and site (both as fixed factors) and by relative hatching date (the difference from the mean laying date in each year and at each site separately) and brood size (both as covariates). We also included the interactions between year and site, brood size and relative hatching date, respectively.

The temperature during the late nestling stage differed substantially between the two years. Mean ± SD of mean daily, maximum and minimum temperature during the trapping periods in the late nestling stage in the two years were; mean daily temperature (°C) 2008: 18.2 ± 0.16; 2009: 14.1 ± 1.57; maximum temperature (°C) 2008: 25.8 ± 0.26; 2009: 19.7 ± 1.95; minimum temperature (°C) 2008: 11.2 ± 0.42; 2009: 9.0 ± 1.15, all differences between years being highly significant (t-test: all: t_89_ > 10.8; p < 0.001). Thus, to evaluate how much of a potential year effect that could be explained by temperature in the final model, we exchanged year with average mean daily, maximum and minimum temperatures, respectively, in three different models. We then evaluated the four models’ fit (including the one with year) to the data with AIC.

For fixed factors with three levels (stage and site), we used post-hoc tests to evaluate differences between groups. When interactions were non-significant, they were removed from the model. Otherwise, full models are presented.

As the response variable is counts, we compared all models using either the Poisson or the Negative Binomial distributions. All models performed much better with the Negative Binomial distribution as judged by their AIC score (all Δ AIC > 2000). This is also corroborated by the fact that the variance in number of midges caught are much larger than the mean (overdispersion), a condition when negative binomial models are recommended (Gardner et al. 1995). Furthermore, the negative binomial regression analyses do not seem to be zero-inflated as the standardized deviance residuals are generally between -2 and 2. Thus, all analyses with no. of caught midges as the dependent variable, were performed as generalized linear model with a negative binomial distribution and a log function.

## RESULTS

Bycatches in the traps (other than *Culicoides* midges) were only recorded in 2009. While the vast majority (>93%) of captured insects in the traps belonged to the genus *Culicoides* (by visual determination) some traps (10% of traps) contained the odd individuals of mosquito (Culicidae), black fly (Simulidae), bird flea (Ceratophyllidae), and beetle (Coleoptera). Mites (Arachnida) was also captured in approximately 10% of traps. A total of 17 out of the 30 sequenced individual midges were successfully identified to species level. All of these were found to belong to the biting midge group (*Culicoides spp*). Of the 13 failed, 11 did not amplify using either of the primer pairs, 2 where sequenced and the obtained sequence matched *Culicoides* sequences but with a sequence difference >10% giving them the identification of *Culicoides spp*. Out of the 17 successfully identified sequences, 2 were identified as *Culicoides segnis*, 7 as *Culicoides kibunensis,* and 8 as *Culicoides pictipennis*.

Of the 19 traps placed in empty nest boxes in 2008, none caught any biting midges, thus data from these boxes are not included in any further analyses. Variation in no. of caught midges in occupied nest boxes was significantly explained by breeding stage (ꭓ^2^ = 61.7; P < 0.001; Fig. 1) whereas site (ꭓ^2^ = 1.02; P = 0.60) and the interaction between stage and site was non-significant (χ^2^_4_ = 4.47; P = 0.35). We found significantly more midges during the 4-day sampling period in boxes during the late nestling stage (mean = 7.33; SE = 2.79) than during both the incubation (Mean = 0.41; SE = 0.18; z.ratio = -6.61; P < 0.001) and early nestling stages (mean = 0.47; SE = 0.20; z.ratio = -6.56; P < 0.001). The incubation and early nestling stage did not differ significantly (z.ratio = -0.29; P = 0.95; Fig. 1).

**Figure 1.**
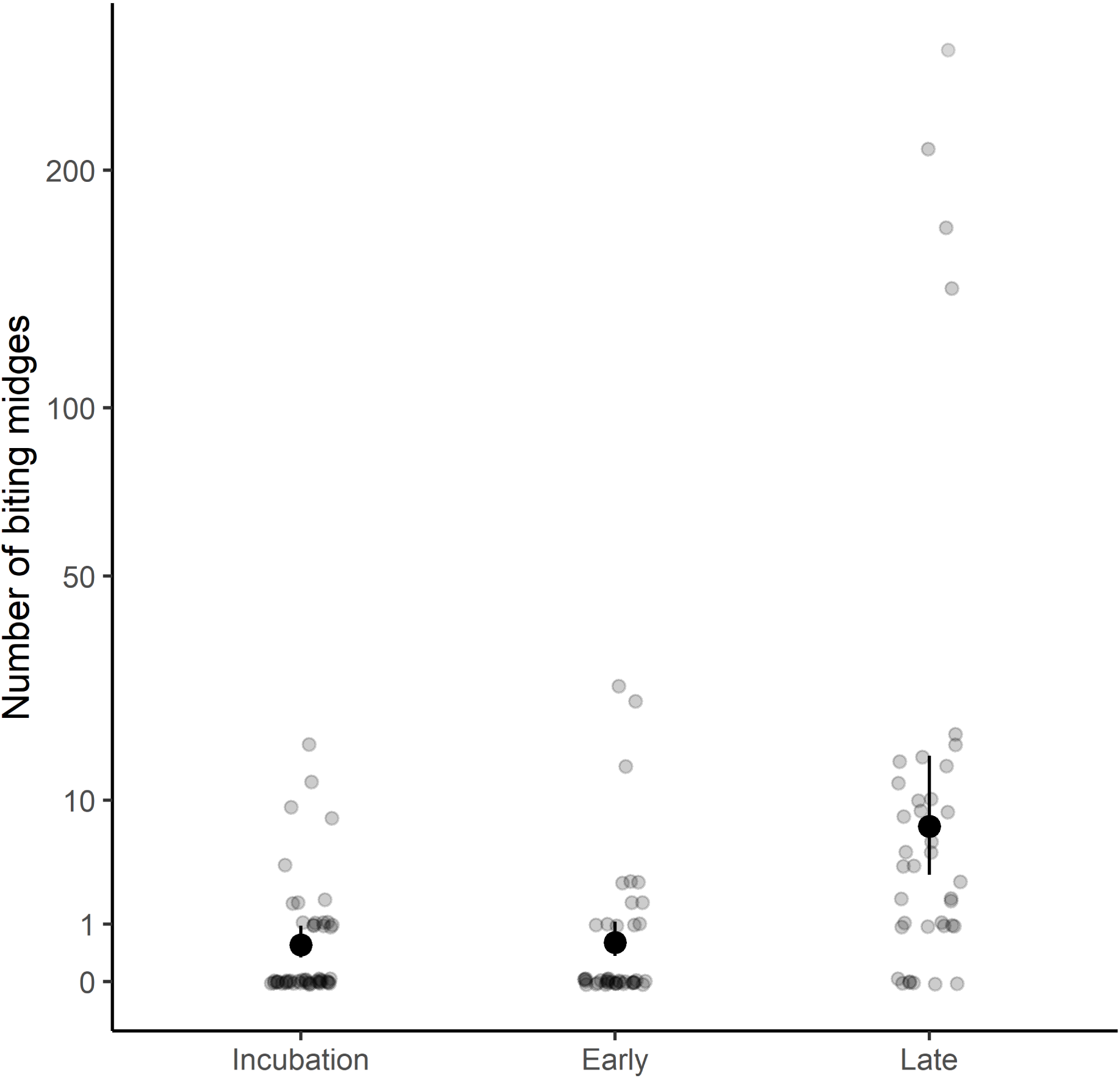
Model estimates of number of trapped biting midges during a 4-day trapping session in 2008 during the incubation stage, the early nestling stage (nestling age 2-6 days) and the late nestling stage (nestling age 12-16 days) in nest boxes occupied by blue tits. Black dots represent the marginal means and bars represent the standard error (SE).

To further investigate which factors could predict the no. of caught midges in the late nestling stage, we combined the data from 2008 and 2009. None of the interactions between year and site, brood size or relative hatching date, respectively, were significant (all ꭓ^2^ < 1.35; P > 0.42). Neither did relative hatching date affect the no. of caught midges during the late nestling stage (ꭓ^2^_1_ = 0.57; P = 0.45). However, brood size positively influenced the abundance of midges in the boxes (ꭓ^2^ = 9.14; P = 0.0025). Furthermore, both year (ꭓ^2^ = 12.3; P = 0.00046) and site (ꭓ^2^ = 14.5; P = 0.00072) explained significant proportions of the variation in no. of caught midges. The overall no. of caught midges per 4-day sampling period in the late nestling stage was larger during 2008 (mean = 10.82; SE = 3.480) than during 2009 (mean = 2.24; SE = 0.614).

We reran the final model of no. of caught midges during the late nestling stage with year exchanged for the average mean, maximum and minimum ambient temperature, respectively. The models including different aspects of ambient temperature had a better fit than the model including year (in comparison to a model with; maximum temperature ΔAIC = 1.7, average mean temperature ΔAIC = 2.2, average minimum temperature ΔAIC = 7.5). Thus, especially minimum ambient temperature seems to be a better predictor of the biting midge abundances within the nest boxes than year as well as compared to the other aspects of ambient temperature. Thus, the number of caught midges increased with average minimum temperature (Fig. 2) and brood size (Fig. 3) but in this model the number of caught midges also increased with relative hatching date (Fig. 4; Table 1). As in the model with year, the number of caught midges differed between sites (Fig. 5) with Revinge nest boxes having more midges than Romele (z.ratio = 3.01; P = 0.0073) and Öved (z.ratio = 3.82; P = 0.0004) whereas Rommele and Öved did not differ (z.ratio = -0.70; P = 0.76).

**Figure 2.**
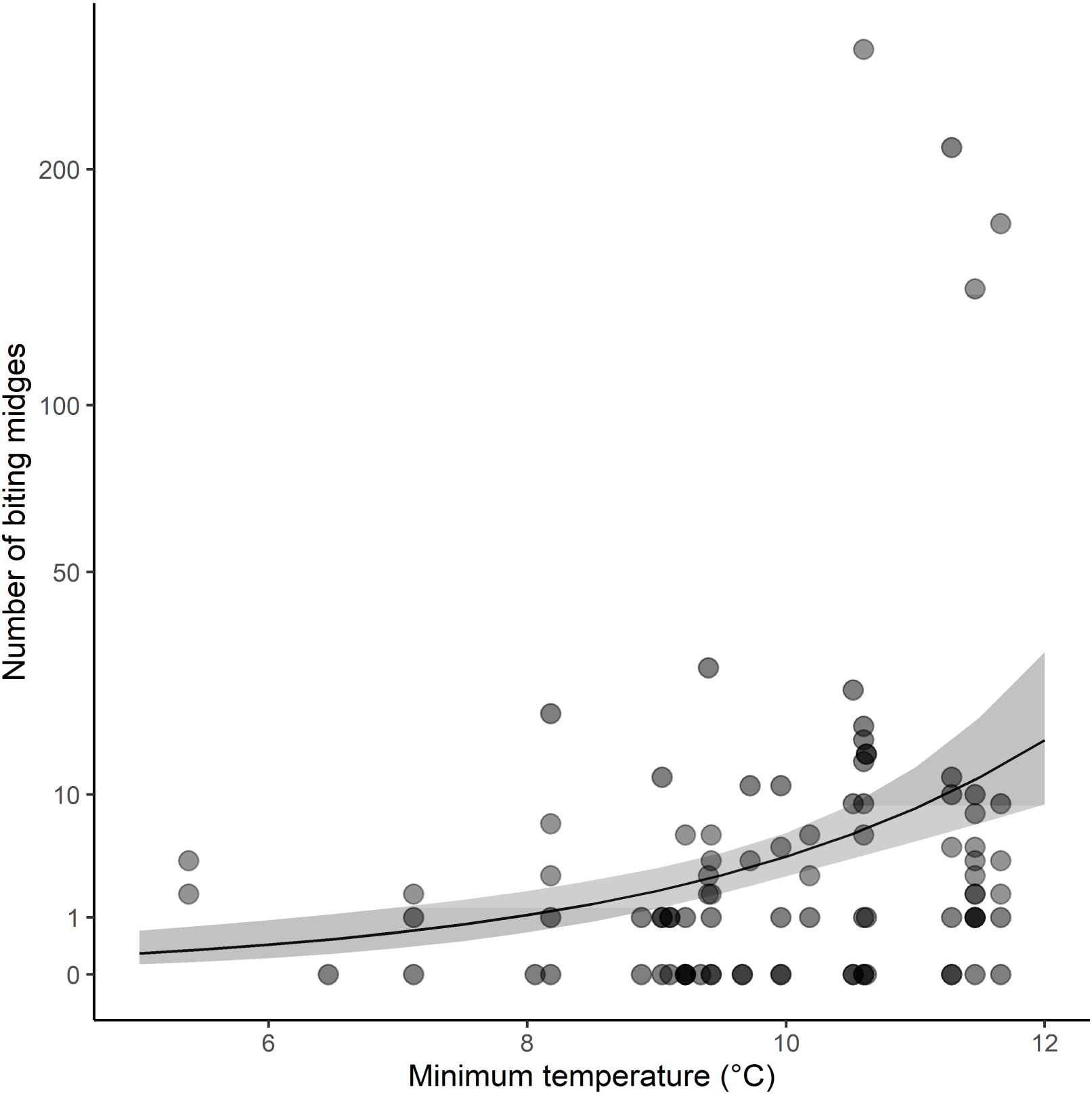
Model estimates of number of trapped biting midges during a 4-day trapping session in 2008 and 2009 during the late blue tit nestling stage (nestling age 12-16 days) in relation to average minimum ambient temperature during the trapping session. Model regression is shown together with their 95 % CI.

**Figure 3.**
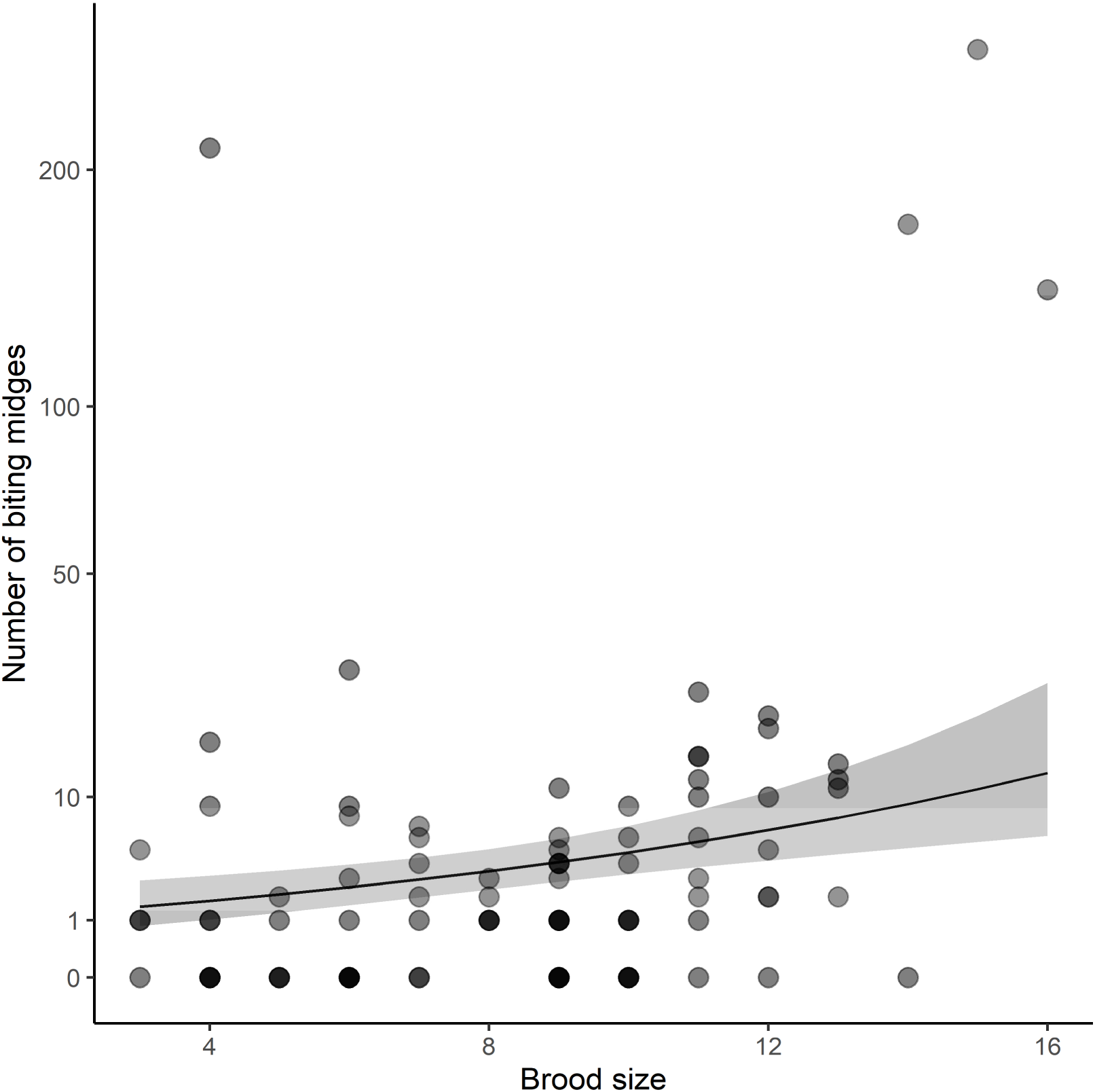
Model estimates of number of trapped biting midges during a 4-day trapping session in 2008 and 2009 during the late blue tit nestling stage (nestling age 12-16 days) in relation to brood size during the trapping session. Model regression is shown together with their 95 % CI.

**Figure 4.**
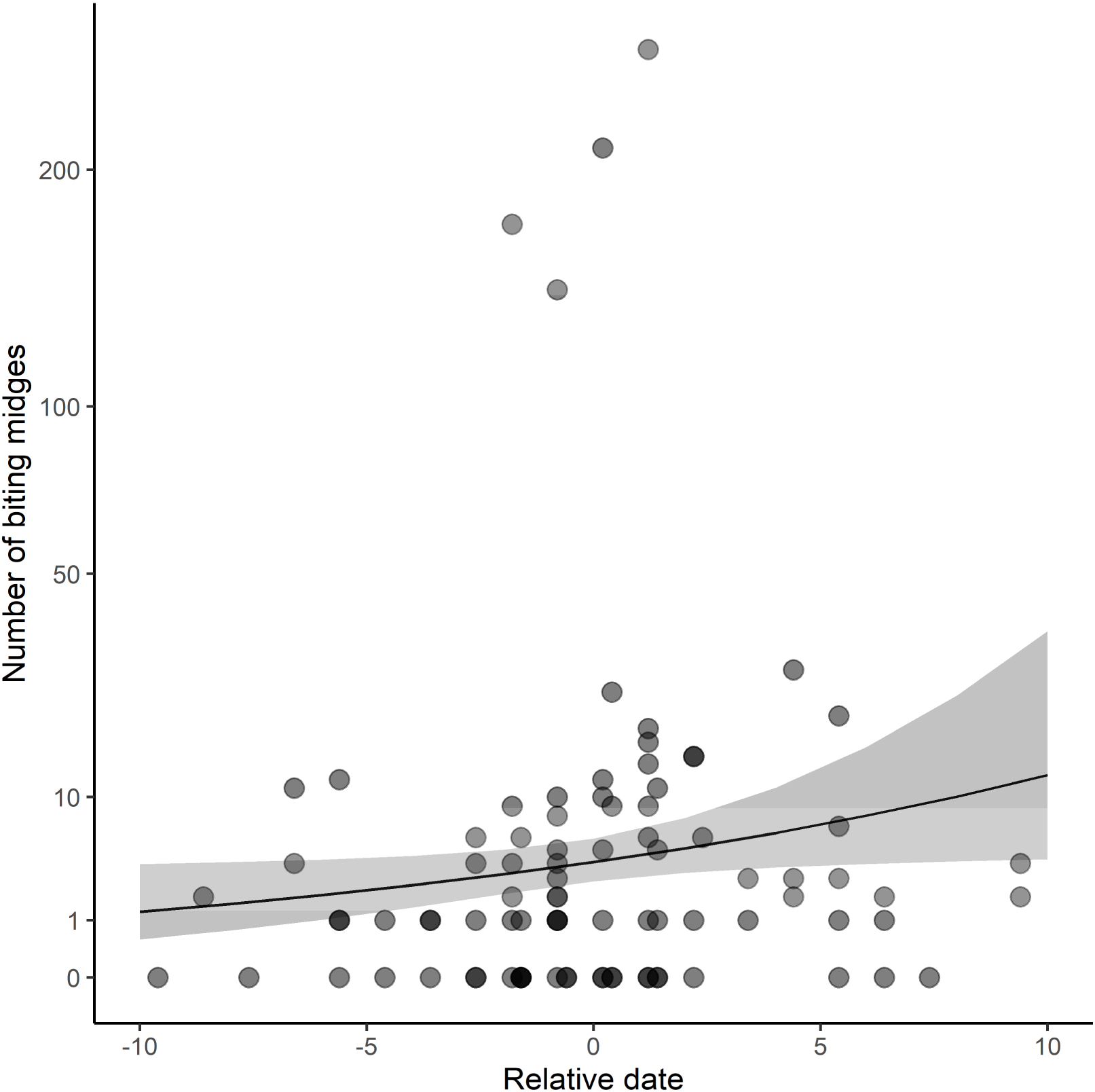
Model estimates of number of trapped biting midges during a 4-day trapping session in 2008 and 2009 during the late blue tit nestling stage (nestling age 12-16 days) in relation to relative hatching date (the difference from the mean laying date in each year and at each site separately). Model regression is shown together with their 95 % CI.

**Figure 5.**
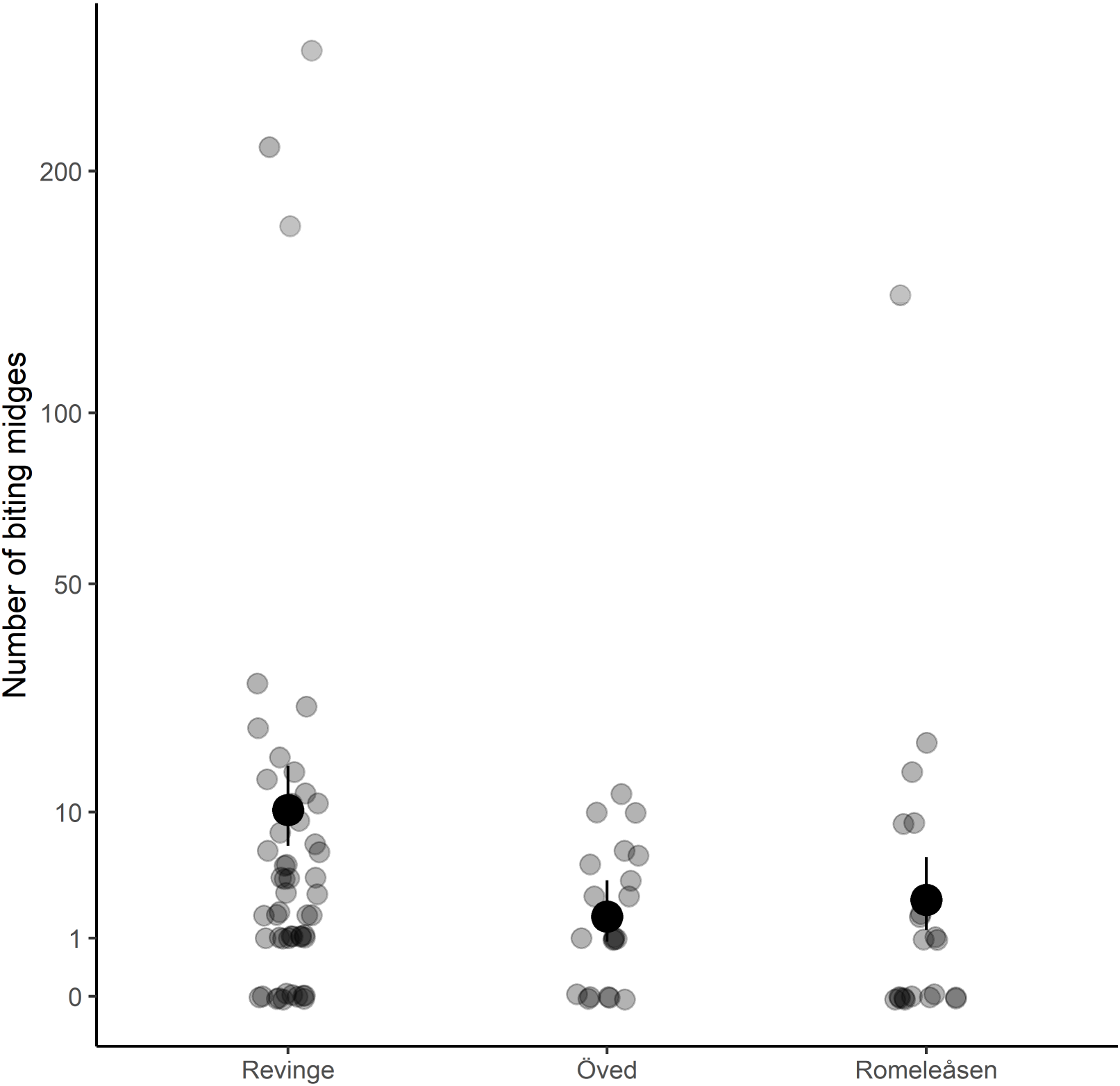
Model estimates of number of trapped biting midges during a 4-day trapping session in 2008 and 2009 during the late blue tit nestling stage (nestling age 12-16 days) at three different sites (Revinge, Öved and Romeleåsen, respectively). Black dots represent the marginal means and bars represent the standard error (SE).

**Table 1.** The influence of brood size, relative hatching date, average minimum ambient temperature and site (Reving, Romele and Öved) on the no. of caught midges per 4-day sampling session in traps placed inside nest boxes occupied by blue tits during the late nestling stage (nestling age 12-16 days) in 2008 and 2009.

| Variable | Estimate (SE) | df | $\chi^2$ | P |
| --- | --- | --- | --- | --- |
| Brood size | 0.16 (0.045) | 1 | 13.07 | 0.00030 |
| Rel. hatching date | 0.11 (0.052) | 1 | 4.70 | 0.030 |
| Min. temperature | 0.688 (0.142) | 1 | 23.56 | <0.00001 |
| **Site** |  | 2 | 19.06 | <0.00001 |
| Site = Revinge | 10.20 (2.22) |  |  |  |
| Site = Romele | 2.74 (1.03) |  |  |  |
| Site = Öved | 1.87 (0.716) |  |  |  |

## DISCUSSION

The vast majority of flying insects caught in our nest box traps belonged to the genus *Culicoides* (biting midges). Commonly biting midges constitute the majority of flies captured in boxes, but black flies (*Simuliidae*) have also been captured with similar traps (Tomás et al. 2008b; Martínez-de la Puente et al. 2009a;b; Castaño-Vázquez et al. 2020) although not always (Votýpka et al. 2009). These results suggest that *Culicoides* midges actively seek out and enter the nest boxes.

All three biting midge species found inside our blue tit nest boxes (*Culicoides segnis*, *C. kibunensis* and *C. pictipennis*) have been found within nest boxes in other studies (Martínez-de la Puente et al. 2009b; Žiegytė et al. 2021). These species prefer birds over other vertebrate hosts as blood source for egg production (Bobeva et al. 2015; González et al. 2022) and have also been verified to have parasite sporozoites in their salivary glands (Chagas et al. 2024), thereby capable of acting as important vectors for *Haemoproteus*.

### Factors affecting biting midge abundances at a general level

Average ambient temperatures during spring have been found to positively affect biting midge abundances in nest boxes (Castaño-Vázquez & Merino 2022 but see Martínez-de la Puente et al. 2009a). As ambient temperatures are generally increasing, the abundances of biting midges are also increasing over time (Castaño-Vázquez & Merino 2022) suggesting an increasing number of *Haemoproteus* infected birds in recent years. Since our study only includes two years, we cannot evaluate any long-term relations to climatic factors but note that the year with high abundances of biting midges (2008) also was much warmer than 2009. Especially, the average minimum ambient temperature proved to have a much better fit to the data than other aspects of ambient temperature or year. Since biting midges are crepuscular or nocturnal (Mellor et al. 2000) and have been found to have reduced activity around the minimum temperatures reported here (Mellor et al. 2000; Tugwell et al. 2021), it seems reasonable that the minimum temperature is affecting their abundance the most.

The abundance of midges within nest boxes varied notably between years being nearly five times higher in 2008 than in 2009. Between-year variation in biting midge abundances has generally been observed (Žiegytė et al. 2021; Castaño-Vázquez & Merino. 2022), indicating that year-specific climatic variation may be an important determinate of midge abundances. Thus, the risk for nestlings to be infected may be highly dependent on in which year they are born. Considering potentially detrimental effects of malaria parasites (Grabow et al. 2025), such cohort effects may explain variation in recruitment rate and overall density between years.

### Factors affecting biting midge abundances at the nest box level

Since we did not find any biting midges in empty boxes (see Tomás et al. (2008a) and Žiegytė et al. (2021) for similar results), cues in relation to the brood seem to attract the midges. Several cues that potentially can be used by biting midges to locate a brood have been suggested, each of them relevant at different ranges (Marzal et al. 2022). At ranges up to 10 m, the broods’ production of CO_2_ might be perceived by the midges. The strength of CO_2_ as an attractant of biting midges relates to how much CO_2_ the brood produces (Castaño-Vázquez et al. 2020). At semi-long ranges, midges might be able to use parental behaviours as guidance to the brood. The feeding rate of blue tit parents has been found to be positively related to the number of captured biting midges within that nest box (Tomás et al 2008b). At very close ranges, the heat produced by the brood might also serve as a cue for midges. Nest temperature is positively related to biting midge abundance in nests of pied flycatchers (*Ficedula hypoleuca*) (Martínez-de la Puente et al. 2010b) and experimentally heated nests resulted in higher infection rate of *Haemoproteus* at a temperate site but not at a Mediterranean one (Castaño-Vázquez et al. 2021). Our findings of an increasing abundance of biting midges with breeding stage and brood size, suggest that total metabolic activity within the nest box will guide the midges. Many nestlings and those of higher age will produce both more CO_2_ (Castaño-Vázquez et al. 2020) and more heat (Nilsson & Nord 2017) making it difficult to separate these two attractants. Furthermore, if warmer springs will induce metabolically costly cooling behavious among nestlings (cf. Andreasson et al. 2018), host-vector encounters can also be predicted to increase in a warmer climate. Since biting midges are crepuscular or nocturnal (Mellor et al. 2000), cues based on parental behaviours (Tomás et al. 2008b) seem to be less relevant as parents decrease their feeding rate towards the evening (Andreasson et al. 2020).

### Factors affecting biting midge abundances at the spatial level

We found pronounced differences between the sub-populations sampled in our study. The area with the highest abundance of biting midges (Revinge) is also the plot with the most standing water, harboring a lake and several ponds and bogs. It is also the area at the lowest altitude of the three study plots. If the Revinge site harbor more biting midges due to on average being moister or some other feature favoring the vectors, these same cues could serve as habitat cues for prospecting breeding blue tits for minimizing host-vector encounters. The extent to which vector abundance vary also within the sites have to await more years of sampling. With the increasing risk of being infected in a warming climate, selection might act on breeding birds to re-evaluate the relative importance of different quality cues when deciding where to breed. Furthermore, the population-level parasite pool could change between years due to variation in immigration rates from adjacent sub-populations.

### Factors affecting biting midge abundances at the temporal level

Whitin a year, the abundance of biting midges are generally increasing with time of the breeding season (Martínez-de la Puente et al. 2009a; Castaño-Vázquez & Merino 2022). Also in our study, the abundance increased with hatching date, at least when minimum temperature was also included in the model (Table 1). Thus, the risk of losing blood and getting infected by *Haemoproteus* increases the later you hatch which can be part of the explanation for generally higher recruitment of early hatched birds (Svensson 1997).

The sometimes high numbers of biting midges found within nest boxes, suggest that at least the *Haemoproteus* parasite is transmitted already to nestlings. As the blood removal by the midges and/or the effect of the parasite infection during the establishment phase may be detrimental to the development of nestlings (Tomás et al. 2008b; Martínez-de la Puente et al. 2010b; Castaño-Vázquez & Merino 2022), parents should be selected to minimise the encounter rate with vectors. This can potentially be achieved by taking vector abundance into account when choosing breeding time and site. Since this cue is potentially becoming more and more important as climate is warming, other important cues may have to be deprioritised compared to past conditions. It is harder to see how parents could avoid the close range attracting cues (mainly CO_2_ and heat production by the brood) although a reduction in brood size would reduce the encounter rate with biting midges. Furthermore, the annual variation in biting midge abundances may have cohort effects with individuals born during springs with relatively low midge abundances being better off.

